# Improving the Reliability and Accuracy of Population Receptive Field Measures Using a Logarithmically Warped Stimulus

**DOI:** 10.1101/2024.06.11.598529

**Authors:** Kelly Chang, Ione Fine, Geoffrey M. Boynton

**Author notes:** Corresponding Author: Kelly Chang.

## Abstract

The population receptive field method, which measures the region in visual space that elicits a BOLD signal in a voxel in retinotopic cortex, is a powerful tool for investigating the functional organization of human visual cortex with fMRI (Dumoulin & Wandell, 2008). However, recent work has shown that population receptive field (pRF) estimates for early retinotopic visual areas can be biased and unreliable, especially for voxels representing the fovea. Here, we show that a ‘log-bar’ stimulus that is logarithmically warped along the eccentricity dimension produces more reliable estimates of pRF size and location than the traditional moving bar stimulus. The log-bar stimulus was better able to identify pRFs near the foveal representation, and pRFs were smaller in size, consistent with simulation estimates of receptive field sizes in the fovea.

## Introduction

The population receptive field (pRF) method is a powerful tool for investigating the functional organization of human sensory cortex with fMRI (Dumoulin & Wandell, 2008). When applied to visual cortex, the region of visual space that elicits a response in an fMRI voxel is estimated by measuring the BOLD activity generated by slowly moving bars of flickering checkerboards and finding the location and size of the Gaussian in visual space that, after accounting for hemodynamic delay, best-predicts the time-course of each voxel. This pRF is thought to represent the collective response of the many thousands of neurons in that voxel. The pRF method has been used in various ways, such as examining spatial attention dynamics (Kay et al., 2015; Klein et al., 2014; Sprague & Serences, 2013), changes due to neural plasticity (Baseler et al., 2011; Haak et al., 2012; Huber et al., 2019), and assessing clinical diagnoses (Barbot et al., 2021; Brewer & Barton, 2012; Schwarzkopf et al., 2014).

The reliability and validity of pRF estimates are still a matter of active research. Previous work has shown that pRF polar angle and eccentricity estimates are, in general, highly reliable (Benson et al., 2018; Senden et al., 2014; van Dijk et al., 2016; Zeidman et al., 2018). However, pRF size estimates vary dramatically across different stimulus configurations (Alvarez et al., 2015; Binda et al., 2013; Infanti & Schwarzkopf, 2020; Linhardt et al., 2021) and model fitting procedures (Lage-Castellanos et al., 2020; Zeidman et al., 2018). pRF size estimates located in foveal regions are particularly sensitive to differences in protocol. Moreover, when using traditional techniques, pRF estimates for the fovea generally have poor model fits and are substantially less reliable (Linhardt et al., 2021; Schira et al., 2007).

The unreliability of some pRF size estimates seems to be related to the large cortical magnification factor and small pRF sizes near the fovea (Alvarez et al., 2015; Linhardt et al., 2021). The cortical magnification factor (CMF) is the amount of visual cortex devoted to a fixed area of visual space (Daniel & Whitteridge, 1961). In all retinotopic visual areas, CMF decreases from the fovea to the periphery. Also, pRF sizes vary inversely with CMF, such that *CMF* ≈ *k*/*σ*_*pRF*_: so that the smallest pRFs are found near the fovea where the CMF is largest (Harvey & Dumoulin, 2011). Roughly 7 times as much cortex represents the central 1 degree of visual space as a 1-degree patch of cortex at 12 degrees in the periphery.

For a given fMRI voxel, the optimal speed and width of a moving bar stimulus depend on the size of the pRF in that location. If the bar moves too quickly, it will sweep through a voxel’s pRF too quickly and produce little or no response due to the slow hemodynamics (blurred over 3-5s) and discrete temporal sampling (typically 1 to 2 seconds) of fMRI. The difficulty with using a very slow bar is that bars that move slowly will take a prohibitively long time to sample the entire visual field. Since cortical magnification and pRF sizes vary throughout the visual field, a bar of fixed width and speed will sweep through pRFs approximately 7 times faster in the fovea than in the periphery. Thus, using a bar of constant width and speed cannot be optimal for the entire visual field.

One method to address this issue has been to combine various speeds and widths that are optimized for both foveal and peripheral pRFs respectively (Benson et al., 2012; Linhardt et al., 2021). This method successfully estimates smaller pRFs near the fovea. However, these methods require long acquisition times because of the need to acquire data for multiple bar configurations. A second approach has been to design multifocal stimuli scaled to cortical magnification – resembling a black and white dartboard with segments pseudorandomly alternating between black and white (Binda et al., 2013). Although the multifocal stimulus successfully produces small pRF estimates, the significantly reduced signal-to-noise for multifocal stimuli greatly increases the acquisition time needed to obtain reliable estimates (Binda et al., 2013; Vanni et al., 2005). Finally, Alvarez et al. (2015) compared two CMF-scaled stimuli: (1) a combined wedge and ring and (2) a logarithmically-scaled bar stimulus. The combined wedge and expanding/contract ring produced pRFs similar in size to the fixed-width drifting bar stimulus. The logarithmic bar produced smaller pRFs, but again suffered from low signal-to-noise. This may be because the entire bar width was scaled based on the bar’s most central location, so the bar region that extended into the periphery was optimized for more foveal regions.

Here, we introduce a novel ‘log-bar’ stimulus: a moving bar logarithmically warped along the eccentricity dimension. This results in a slowly-traveling, thin bar near the fovea with a faster-traveling wider bar out in the periphery. Because of the logarithmic nature of the mapping from visual space to cortex (Schwartz, 1980), the log-bar stimulus is designed to approximate a constantly moving fixed-sized bar on the cortical surface, whose speed and size can be optimized across the entire visual field simultaneously.

Finally, using a simple cortical model, we closely replicated fMRI results for both fixed- and log-bar stimuli, including the finding that the log-bar recovers pRF parameters more accurately and reliably than a fixed-sized bar stimulus, especially for more foveal pRFs.

## Methods

### Participants

Data were acquired from 12 participants (6 females, 6 males), ages 20 – 38 (*M* = 28.67, *SD* = 5.66). All participants were paid and had normal or corrected-to-normal acuity. All gave informed consent in accordance with the human participant Institutional Review Board at the University of Washington, in adherence with the Declaration of Helsinki.

### MRI Acquisition

MRI data were acquired at the University of Washington Center for Human Neuroscience (CHN) on a Siemens 3T Prisma with a 64-channel phase-array head coil.

The protocol included 3D volumetric navigator-guided T1-weighted (T1w, multiecho) and T2-weighted (T2w) images acquired at 0.8 x 0.8 x 0.8 mm isotropic resolution (FOV = 166.4 x 240 x 256 mm3, matrix size = 208 x 300 x 320; Tisdall et al., 2012).

BOLD data were acquired using a T2*-weighted 2 x 2 x 2 mm isotropic multiband gradient-echo (multiband acceleration = 4), echo-planar imaging sequences (TR / TE = 1200/30 ms, flip angle = 64°, FOV = 212 × 212, matrix size = 106 × 106, 56 oblique axial slices). Each run collected 305 volumes (approximately 6.1 minutes), and we collected a total of 12 runs with 6 runs per stimulus condition across two sessions for each participant. Additionally, a pair of opposite phase-encoded EPI references were acquired for each session.

### Preprocessing

Data were processed using the fMRIPrep-21.0.1 pipeline (Esteban et al., 2019). The following details are based on fMRIPrep’s documentation. The T1w image was corrected for intensity non-uniformity with N4BiasFieldCorrection (ANTs-2.3.3; Tustison et al., 2010) and used as the T1w-reference throughout the workflow. The T1w-reference was then skull-stripped with a Nipype implementation of the antsBrainExtraction.sh workflow. Brain tissue segmentation of cerebrospinal fluid (CSF), white matter (WM), and gray matter (GM) was performed on the brain extracted T1w using FSL’s fast segmentation tool (Zhang et al., 2001). Brain surfaces were reconstructed using FreeSurfer recon-all (Dale et al., 1999).

A B0-nonuniformity map (or field map) was estimated based on two echo-planar imaging (EPI) references with topup from FSL (Andersson et al., 2003). A field map was estimated for each session, and the correction was applied to all functional data within the collected session.

BOLD runs were slice-time corrected to 0.549s (50% of the slice acquisition range 0s – 1.1s) using 3dTshift from AFNI (Cox & Hyde, 1997). The BOLD reference was then co-registered with the T1w reference using bbregister (FreeSurfer), which implements boundary-based registration (Greve & Fischl, 2009). Co-registration was configured as a rigid-body transformation with six degrees of freedom. The BOLD time series was then resampled onto participant-specific surface space in a non-gridded resampling performed using mri_vol2surf (FreeSurfer).

Head-motion parameters with respect to the BOLD acquisitions were estimated and expanded by including temporal derivatives and quadratic terms for each parameter (Satterthwaite et al., 2013). Additionally, a set of physiological regressors was extracted to allow for component-based noise correction (CompCor Behzadi et al., 2007). After high-pass filtering the preprocessed BOLD time series (using a discrete cosine filter with 128s cut-off), principal components were estimated for the anatomical variant (aCompCor). The aCompCor components were calculated separately within the WM and CSF masks.

All BOLD data underwent temporal high-pass filtering (Fourier GLM, cut-off: 3 cycles) and were denoised with 24 head-motion parameters (rigid-body, temporal derivative, quadratic terms; Satterthwaite et al., 2013) and the first 5 aCompCor (WM + CSF) components (Behzadi et al., 2007).

### Stimuli

Stimuli were presented on a BOLDscreen32 LCD monitor (Cambridge Research Systems, Rochester, UK) operating at a resolution of 1920 × 1080 at 60 Hz with a viewing distance of 133 cm. An Intel i7 Mac Mini running MATLAB R2022a (MathWorks Inc., Natick, MA) controlled custom stimulus presentation code based on Psychtoolbox-3 (Brainard, 1997; Pelli, 1997). Behavioral responses were recorded using a button box (Current Designs, Haverford, PA, USA).

Both bar stimuli consisted of an 8 Hz contrast reversing checkerboard texture. Bars were constrained to a central 16° diameter circular aperture. The display was uniform gray outside the aperture.

The experiment consisted of two sessions containing twelve functional runs. In each run, two different types of bars were presented (fixed-bar and log-bar). Each run began with a 2s blank period, followed by eight 45s sweeps of the bar across the full aperture, each with a pseudorandomly chosen direction that was randomly seeded. Each run ended with a 4s blank period, resulting in a total duration of 366s per run. The exact stimuli used in the fixed-bar runs were transformed into log-bar stimuli.

**Figure 1** shows example frames from a single sweep of the fixed-bar and log-bar stimuli used in these experiments. See the OpenNeuro repository for MATLAB/Python code that demonstrates how to warp an image into log-eccentricity space.

**Figure 1.**
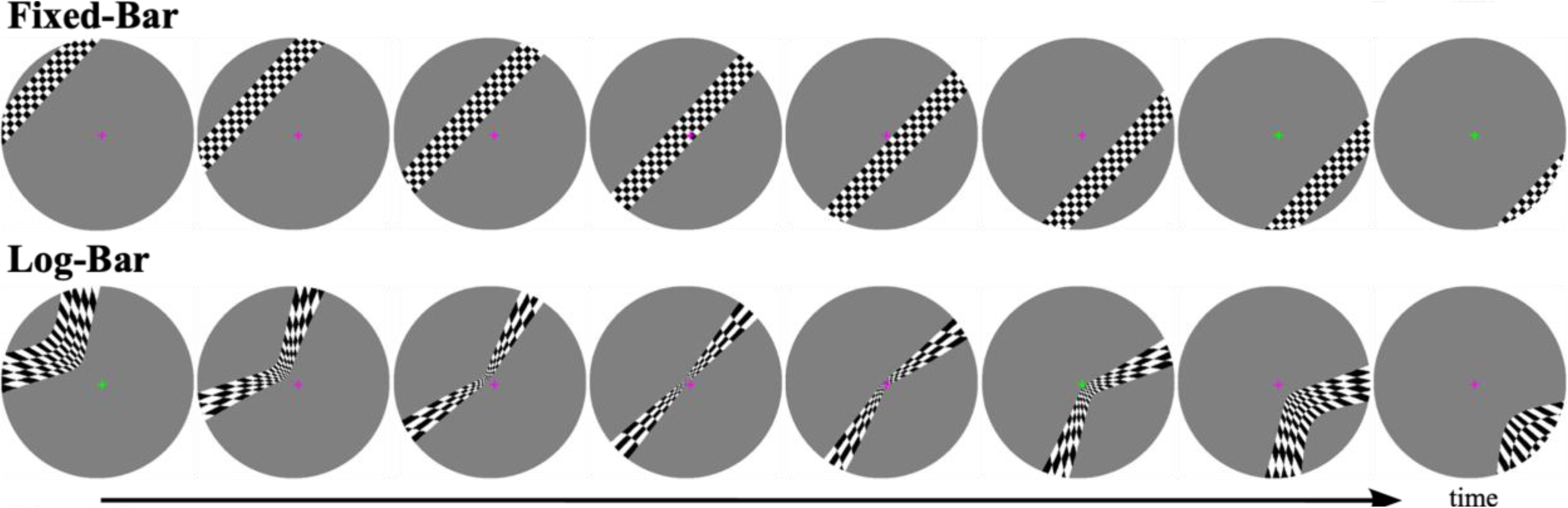
Example frames from the fixed-bar and log-bar stimuli. Log-bar stimuli were produced by distorting the respective fixed-bar frame along the eccentricity axis, as described in Equation 1. Participants were instructed to fixate on the central cross and report with a button press when the cross changed color.

All participants participated in 2 one-hour scanning sessions, with up to a 2-week interval between sessions. Each session consisted of 3 runs of the fixed-bar stimulus and 3 runs of the log-bar stimulus.

### Fixed-bar Stimuli

These stimuli were generated based on the drifting bar from Dumoulin & Wandell (2008). The fixed-bar stimulus was 2° in width and traveled at a constant speed of 0.4 °/s.

### Log-bar Stimuli

To create the log-bar stimuli, each frame of the fixed-bar stimulus was distorted along its eccentricity axis, expressed as:

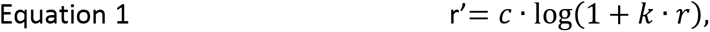

where *r* is the eccentricity (in degrees of visual angle) of each pixel, *k* is a distortion factor, *c* is an overall scale constant, and *r’* is the mapped eccentricity of that pixel for the log-bar stimulus.

Note that this logarithmic warping is consistent with the mapping from visual space to cortical space in the primary visual cortex (Duncan & Boynton, 2003; Schwartz, 1980), which is why the log-bar stimulus produces a wave of activity moving at roughly a constant speed when projected to the cortex.

The distortion factor, *k*, determines the strength of the logarithmic warping. In the limit, as *k* goes to zero, Equation 1 goes toward the identity:

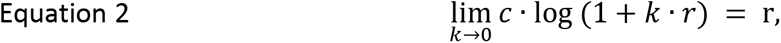

which leads to zero distortion. As *k* increases, more of the original stimulus is warped toward the fovea. For this experiment, our distortion factor was set at *k* = 5, which is roughly consistent with the mapping from visual to cortical space in human visual cortex (Duncan & Boynton, 2003).

The scale factor, *c*, was set as 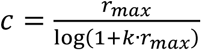, where *r_max_* = 8 (the maximum eccentricity of the stimulus). This choice of *c* ensures that a pixel at 8° in the fixed-bar stimulus is mapped to 8° in the log-bar stimulus.

**Figure 1** shows example frames from a single sweep of the fixed-bar and log-bar stimuli used in these experiments. See the OpenNeuro repository for MATLAB/Python code that demonstrates how to warp an image into log-eccentricity space.

### Experimental Task

A small cross (“+”) was presented at the center of the display and was present continuously throughout the experiment. The color of the cross switched from green to magenta on average 15 times per run, with a minimum inter-switch delay of 3.2s. Participants were instructed to maintain fixation on the cross and to press a response button whenever the color of the cross changed. The purpose of this task was to encourage fixation for the entire scan duration.

### pRF Analysis

#### Region of Interest (ROI) Selection

Initial visual cortex region selection for pRF fitting was performed by aligning each participant’s anatomical surface with Benson’s atlas of the visual cortex using neuropythy (Benson & Winawer, 2018). Areas V1, V2, and V3 were extracted and dilated to ensure the regions of interest (ROI) encompassed the edges of each visual area. This process created an early visual cortex region of interest for each participant and hemisphere used in the pRF fitting process.

Benson’s atlas was used again after pRF fitting to fine-tune the visual area boundaries by including participants’ pRF information from all stimulus types and runs. Finally, manual examination and editing were performed to create final V1, V2, and V3 ROIs for each individual.

Separately, an anatomical definition of the foveal confluence was drawn by dilating a seed placed at the occipital pole to be at least 350 mm^2^ in surface area. The foveal confluence was then functionally constrained to vertices with pRF eccentricity estimates that were ≤ 1.5°. This procedure defined the foveal confluence ROI for each individual. A subset of the vertices in the foveal confluence intersects with vertices in the more foveal regions of V1, V2, or V3.

#### Stimulus Preprocessing

Prior to model fitting, we binarized the stimulus movies, such that 1s indicated the presence of the bar and 0s represented the background. To reduce the computational burden, each frame was spatially linearly downsampled from 540 × 540 pixels to 108 × 108-pixel resolution and temporally linearly downsampled to 366 frames to match the temporal resolution of the fMRI data.

#### Hemodynamic Response Function (HRF)

We modeled each participant’s individual hemodynamic response function using SPM’s gamma function (Friston et al., 1998). The HRF is characterized by 6 parameters expressed as:

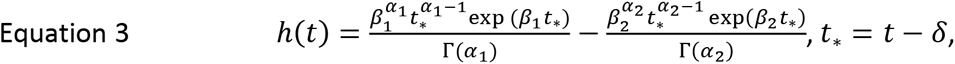

where *δ* is the onset delay in seconds, *⍺*_1_ is the time to peak response in seconds, *⍺*_1_ is the time to undershoot response in seconds, *β*_1_ is the response dispersion, *β*_2_ is the undershoot dispersion, and *c* is the response-to-undershoot ratio. An HRF function was estimated for each hemisphere. This estimation resulted in 12 participants × 2 hemispheres = 24 sets of HRF parameters. There were no significant differences in participant HRF parameters.

#### pRF Modeling

We modeled each vertex as an isometric 2-D Gaussian with 3 parameters of interest. The model function was expressed as

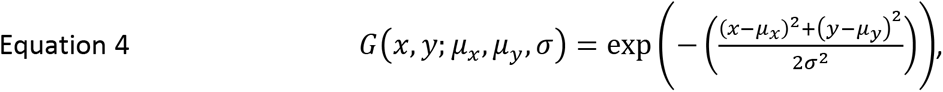

where *μ*_*x*_ and *μ*_*y*_ defined the center and *σ* defined the standard deviation of the pRF 2D Gaussian.

The pRF method assumes that the fMRI time series for each vertex is the dot product of the stimulus time series convolved with each participant’s HRF in time and the vertex’s pRF Gaussian model in space. For each vertex, the pRF model can be formally expressed as:

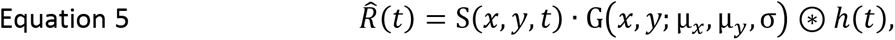

where *R̂*(*t*) is the predicted fMRI time series, *S*(*x*, *y*, *t*) is the stimulus movie, *G*(*x*, *y*; *μ*_*x*_, *μ*_*y*_, *σ*) is a 2D Gaussian, and *h*(*t*) is the hemodynamic response function.

Using custom MATLAB software, pRF estimates for each vertex were determined to be the values that maximized the correlation between the predicted and actual fMRI time course. The square of this correlation is the proportion of variance in the fMRI time series explained by the pRF model. The pRF estimation yields three parameters: pRF center location (*μ*_*x*_, *μ*_*y*_) and pRF size (*σ*).

We conducted pRF model fitting in two stages. First, a coarse parameter grid-search was performed with seeds for pRF centers (*μ*_*x*_, *μ*_*y*_) linearly sampled from - 8° to 8° in 20 steps and seeds for pRF size (*σ*) linearly spaced from 1 to 5 in 20 steps. The pRF size seeds were chosen to be larger than expected because Lage-Castellanos et al. (2020) reported steeper convergence gradients for pRF size when starting from larger values, irrespective of true pRF size. A predicted time course was generated for each seed and correlated with the actual fMRI time course. The seed parameters with the highest correlation were chosen as the initial starting parameters for the next stage of fitting.

Second, we estimated each participant’s HRF for each hemisphere by holding the pRF parameters from the grid-search fixed and fitting for the 6 HRF (*δ*, *⍺*_1_, *⍺*_2_, *β*_1_, *β*_2_, *c*) parameters. Then, we held the HRF parameters constant and fit the pRF parameters. This process was repeated for three iterations to ensure parameters converged on a stable solution. To limit computation time, the HRF estimation process was carried out by selecting 15% of the vertices out of the subset of vertices with explained variance > 20%. The median HRF parameters of these fitted voxels were used to estimate that individual’s HRF. This HRF fitting procedure used all collected fMRI data for each participant, irrespective of session and stimulus type, to prevent biasing the final pRF estimates towards any particular stimulus type or session.

After participant HRFs were estimated, the final stage of pRF estimation was performed. The best-starting seed parameters were once again calculated from the coarse grid-search procedure, and then a nonlinear minimization routine (MATLAB’s fminsearch) was used to find the final pRF estimates.

pRF estimates were always fit separately for each stimulus type (fixed-bar and log-bar). We estimated these parameters for each session as well as collapsed across both sessions (session 1, session 2, sessions 1+2). This procedure resulted in 12 participants × 2 hemispheres × 2 stimulus types × 3 session configurations = 144 sets of pRF estimates.

After pRF fitting, the Cartesian coordinates of the pRF center (*μ*_*x*_, *μ*_*y*_) were transformed into polar coordinates (polar angle and eccentricity). Only vertices with estimated pRF centers less than 8° eccentricity (i.e., within the display), had a pRF size greater than 0.05°, and whose model variance explained was > 10% were retained for subsequent analyses.

### pRF Model Simulations

To gain a deeper understanding of why differences between fixed- and log-bar stimuli produced different receptive field sizes, we carried out a simulation experiment.

First, we defined a unique set of 4800 pRF locations by sampling polar angle coordinates from 0° to 345° in 15° steps and logarithmically sampling eccentricity coordinates from 0.01° to 8° in 200 steps. pRF size was fixed to be linearly related to eccentricity value, such that *σ* = 0.15 ∗ eccentricity + 0.1, derived from Himmelberg et al. (2021). Each pRF was defined as a Gaussian, as described in Equation 4.

fMRI time courses were then generated for each simulated pRF, for both fixed- and log-bar stimuli, as the dot product of the stimulus aperture with the simulated Gaussian pRF followed by a convolution with a realistic hemodynamic response function. Lastly, IID Gaussian noise was added to the simulated time courses with a standard deviation chosen to match the average variance of our fMRI data. These simulated time courses were generated 100 times to create a simulated dataset that matched our collected data.

The simulated data were fit using the pRF fitting procedures described above. As for the real data, only simulated pRFs with estimated pRF centers less than 8° eccentricity, pRF sizes greater than 0.05°, and variance explained > 10% were retained for all subsequent analyses.

## Results

### Behavioral Task

Participants’ responses to the color detection task were analyzed across 12 functional runs. Performance was defined as:

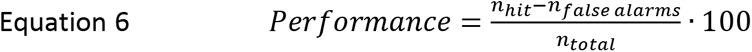

where *n*_*hit*_ is the number of successful color change detections (defined as a button press within 1s of a color change), *n*_*false alarms*_ indicates the number of additional responses without a color change, and *n*_*total*_ indicates the total number of color changes. Average behavioral performance was 96.39%, and the interquartile range of performance values across participants was (95.28%, 98.89%).

### pRF Cortical Maps

For both fixed- and log-bar stimuli, we generated pRF parameter estimates of polar angle, eccentricity, pRF size, and variance explained for all participants. **Figure 2** shows cortical maps of each parameter for an example participant.

**Figure 2.**
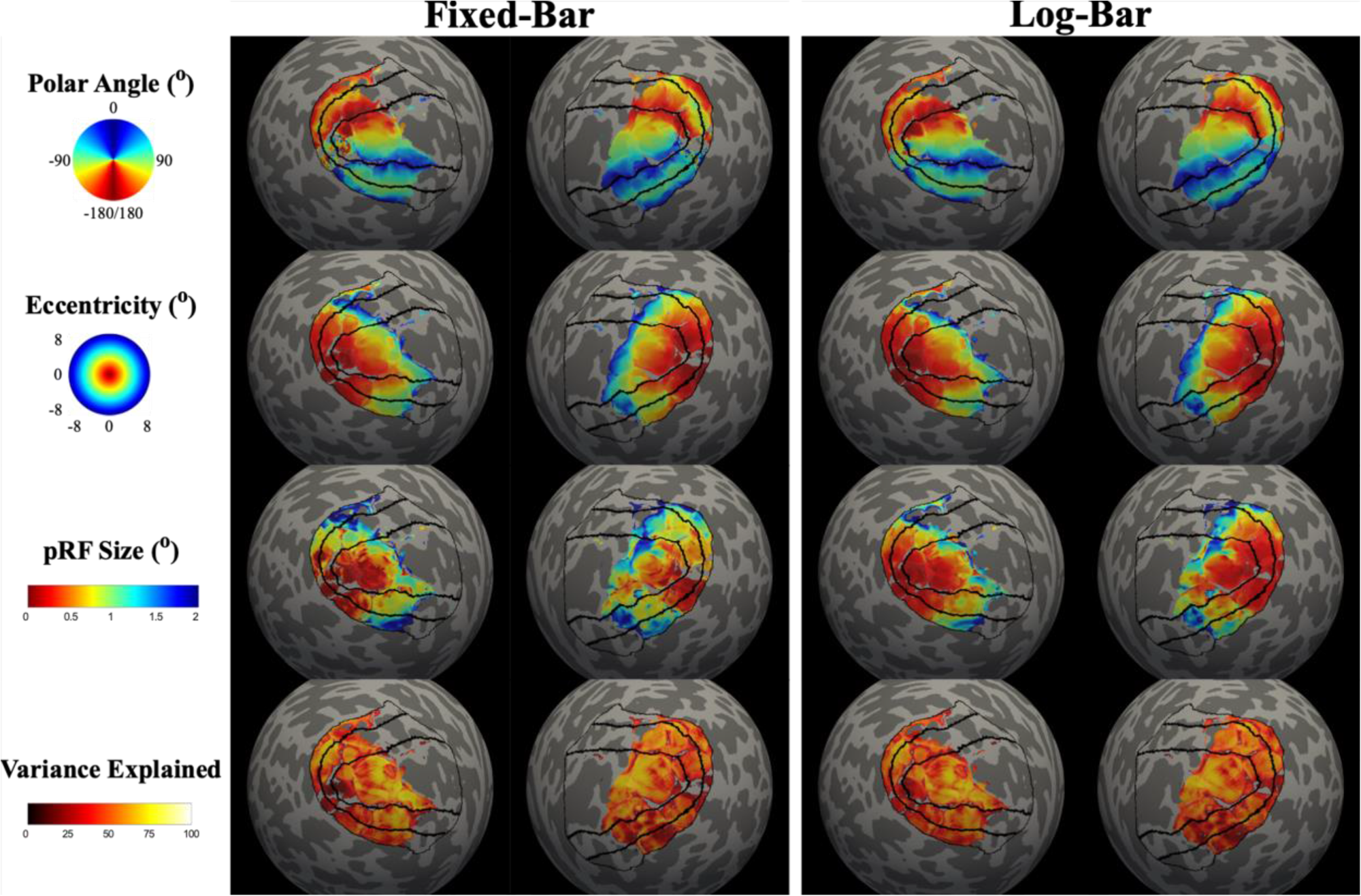
Cortical maps of pRF estimates for an example participant. Solid black lines represent V1, V3, and V3. pRF estimates were thresholded at > 10% variance explained.

### pRF Estimate Intersession Reliability

Next, we compared the intersession reliability of pRF estimates derived from fixed- and log-bar stimuli using the procedure described in (van Dijk et al., 2016). Briefly, Spearman’s rank correlation coefficients were calculated between pRF parameter estimates from session 1 and session 2 of the same stimulus type. For our regression analysis, vertex-by-vertex correlation coefficients (for polar angle is a circular statistic, we used the circular correlation coefficient) for each parameter were transformed into Fisher’s z-scores via the arctanh function that converts the non-normal correlation (*r*) sampling distribution into a standardized statistic:

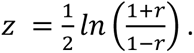

**Figure 3** shows intersession reliability for polar angle, eccentricity, and pRF size estimates for the fixed-bar and log-bar stimulus for the foveal confluence and V1-V3.

**Figure 3.**
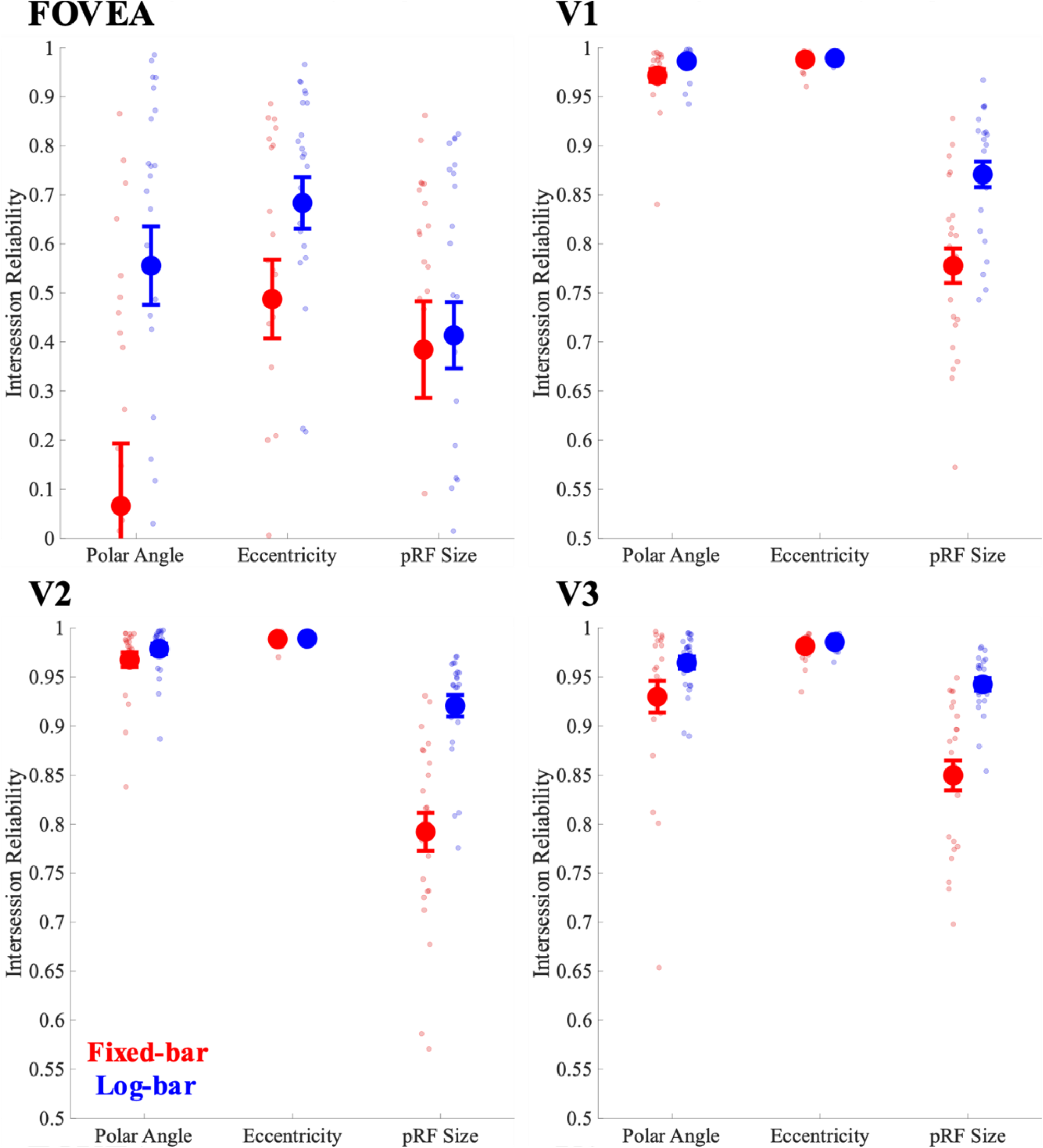
Intersession reliability for pRF parameter estimates of polar angle, eccentricity, and pRF size across visual areas. Red values represent the fixed-bar stimuli and blue values represent the log-bar stimuli.

We compared the intersession reliability of parameter values for fixed- and log-bar stimuli using a mixed-effects ANOVA that treated participants as a random-effect variable, and hemisphere and ROI as fixed-effect variables. There was a main effect of stimulus type (fixed- vs. log-bar) on polar angle (F(1,162) = 34.72, p < 0.0001, η^2^=0.18), eccentricity (*F*(1,162) = 7.34, *p* = 0.0075, η^2^=0.04), and pRF size (*F*(1,162) = 75.35, *p* < 0.0001, η^2^=0.32) such that polar angle, eccentricity, and pRF size estimates were more reliable for the log-bar than the fixed-bar stimulus.

Consistent with previous studies (Alvarez et al., 2015; Benson et al., 2018; van Dijk et al., 2016), there was a main effect of ROI on intersession reliability for polar angle (*F*(3,162) = 173.04, *p* < 0.0001, η^2^=0.76), eccentricity (*F*(3,162) = 390.38, *p* < 0.0001, η^2^=0.88), and pRF size (*F*(3,162) = 129.41, *p* < 0.0001, η^2^=0.71) estimates. A Tukey-Kramer post-hoc analysis revealed that the foveal confluence ROI was consistently the least reliable for all parameters, relative to V1, V2, and V3.

Finally, there was a significant interaction for intersession reliability between ROI and stimulus type for polar angle (*F*(3,162) = 3.06, *p* = 0.03, η^2^=0.05), eccentricity (*F*(3,162) = 3.67, *p* = 0.01, η^2^=0.06), and pRF size (*F*(3,162) = 7.80, *p* < 0.0001, η^2^=0.13). Tukey-Kramer analyses showed that, for polar angle and eccentricity, this effect was driven by lower intersession reliability for the fixed-bar than the log-bar stimulus in the foveal confluence. For pRF size, intersession reliability was lower for the fixed-than the log-bar stimulus in all ROIs except the foveal confluence, where reliability was approximately equal for fixed and log-bar stimuli.

Overall, pRF estimates of polar angle and eccentricity showed very high intersession reliability across all ROIs (foveal confluence, V1, V2, and V3), regardless of whether the fixed-bar or the log-bar stimulus was used. In contrast, the intersession reliability for pRF size was better for the log-bar stimulus than the fixed-bar stimulus in V1, V2, and V3. In the foveal confluence, the log-bar and fixed-bar stimuli showed similar intersession reliability, though far more vertices survived the >10% variance criterion, as described in the next two sections.

### Frequency of Successfully Estimating pRFs

On average, the log-bar stimulus found more vertices that survived the >10% variance explained criterion than the fixed-bar stimulus across all ROIs, as shown in **Figure 4**.

**Figure 4.**
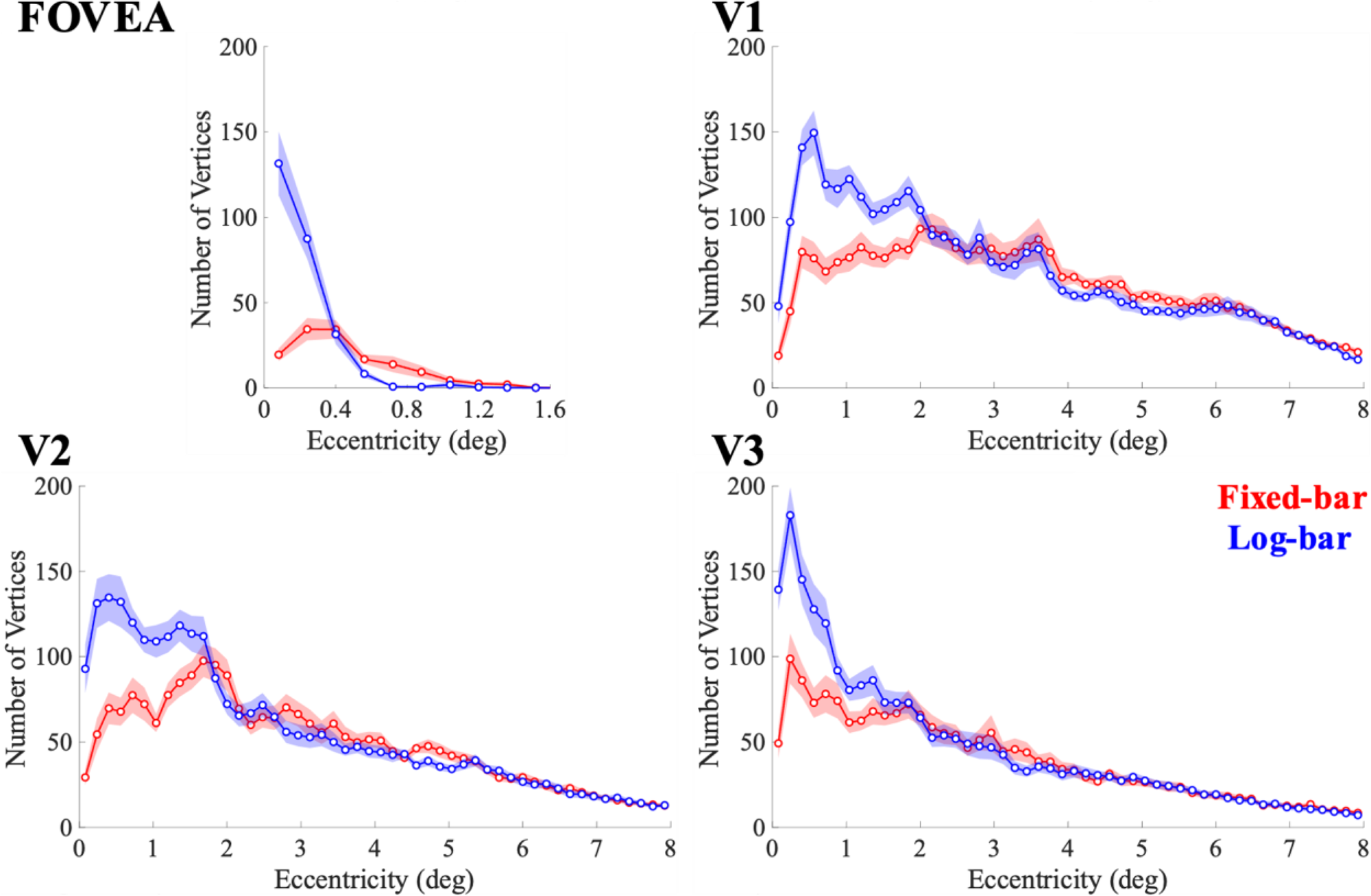
The number of vertices whose pRF fits survived the >10% variance explained criterion as a function of eccentricity. The solid lines represent the average values from the fixed-bar (red) and log-bar (blue) stimuli.

The superior performance of the log-bar stimulus seemed to be confined to the central 2 degrees of visual eccentricity. Within the central 2° of eccentricity, the log-bar stimulus significantly outperforms the fixed-bar. The crossing of the curves in the fovea at 0.4° eccentricity is likely caused by a bias for the fixed-bar stimulus that shifts estimates of foveal pRFs toward more eccentric locations, as described below.

### pRF Parameter Estimates

The pRF parameters were estimated separately for each stimulus aperture, collapsing across sessions, to produce pRF estimates for each participant for each hemisphere.

#### Variance Explained

There was a significant main effect of ROI (*F*(3,77) = 51.60, *p* < 0.0001, η^2^=0.67) on variance explained by the pRF model fitting. Bonferrroni-corrected post-hoc paired t-tests for each ROI 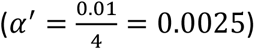 revealed that the variance explained was significantly greater when using the log-bar stimuli within the foveal confluence (*t*(22) = −6.97, *p* < 0.0001, Cohen’s d = 1.45).

#### Eccentricity

**Figure 5** shows pRF eccentricity estimates binned as a function of cortical distance from the foveal center.

**Figure 5.**
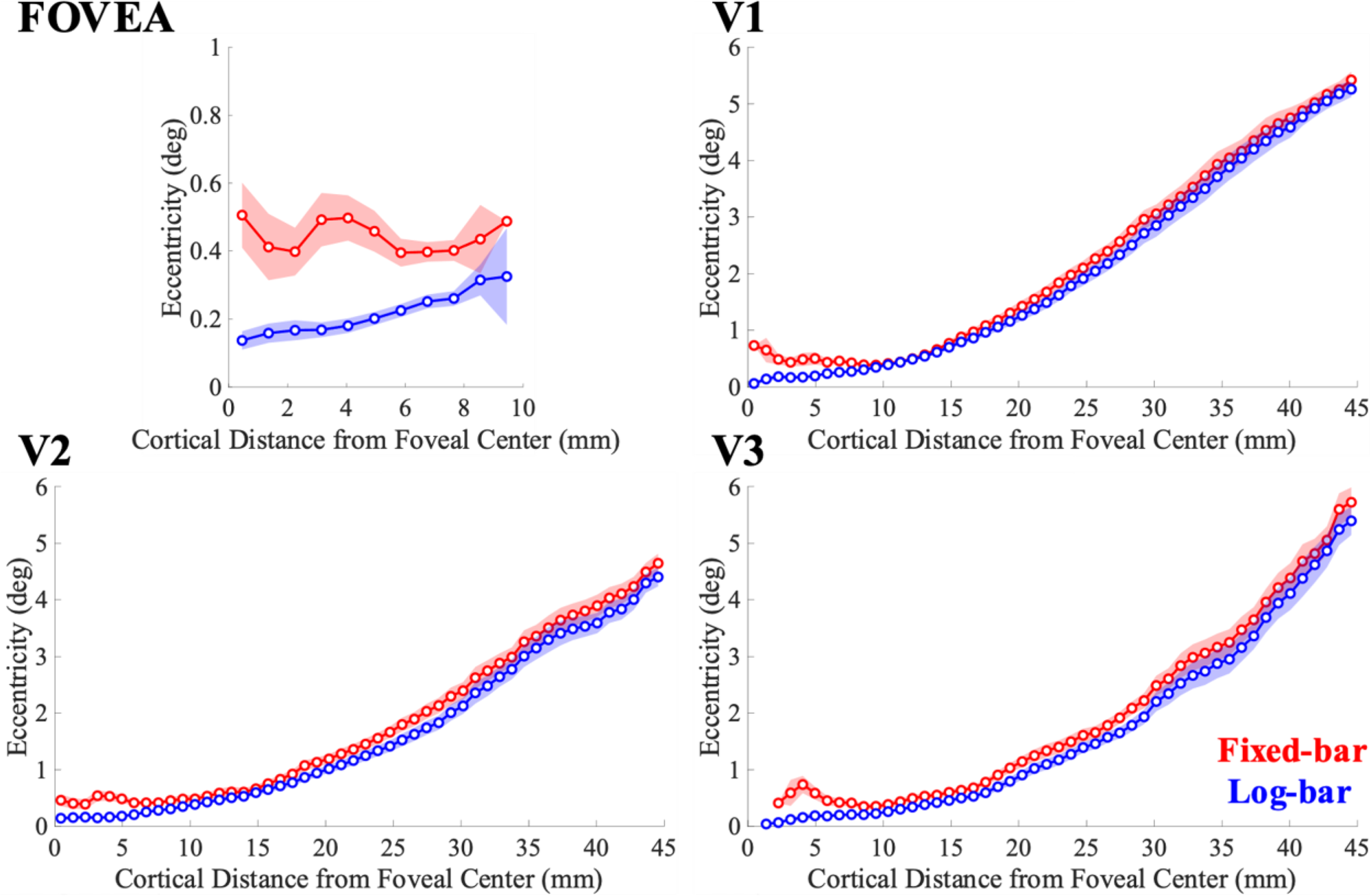
pRF eccentricity estimates binned as a function of cortical distance from the foveal center. Solid lines represent the average across participants for the fixed-bar (red) and log-bar (blue) stimuli and the shaded area represents the ±1 standard error of the mean.

In general, there is a tendency for the fixed-bar stimulus to produce larger estimates of eccentricity, and this tendency is pronounced near the fovea: within the foveal ROI, pRF eccentricity estimates for the fixed-bar stimulus are much larger than for the log-bar stimulus. Indeed, for the fixed bar stimulus, there are few pRFs with eccentricity estimates < 0.4 °: clearly suggesting systematic biases.

We used a repeated-measures ANOVA, with participants as a random-effect variable and hemisphere and ROI as fixed-effect variables, to examine the effects of stimulus type on eccentricity estimates. We found that there was a significant main effect of ROI (*F*(3,76) = 16.48, *p* < 0.0001, η^2^=0.39) on eccentricity differences. Bonferroni-corrected post-hoc paired t-tests 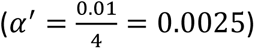 revealed that eccentricity values were significantly larger for the fixed-bar stimulus than the log-bar stimulus in the foveal confluence (*t*(22) = 5.01, p = 0.0001, Cohen’s d = 1.04), V1 (*t*(23) = 3.50, *p =* 0.0019 Cohen’s d = 0.72), V2 (*t*(23) = 6.77, *p* < 0.0001, Cohen’s d = 1.38), and V3 (*t*(23) = 12.17, *p* < 0.0001, Cohen’s d = 2.48).

#### pRF Size

**Figure 6** shows pRF size estimates binned across pRF eccentricity. The fixed-bar stimulus produced larger estimates of pRF size in the foveal confluence. In V1 and V2 there is a ‘cross-over’ near 4° eccentricity, whereas in V3 the log-bar stimulus consistently produces smaller size estimates.

**Figure 6.**
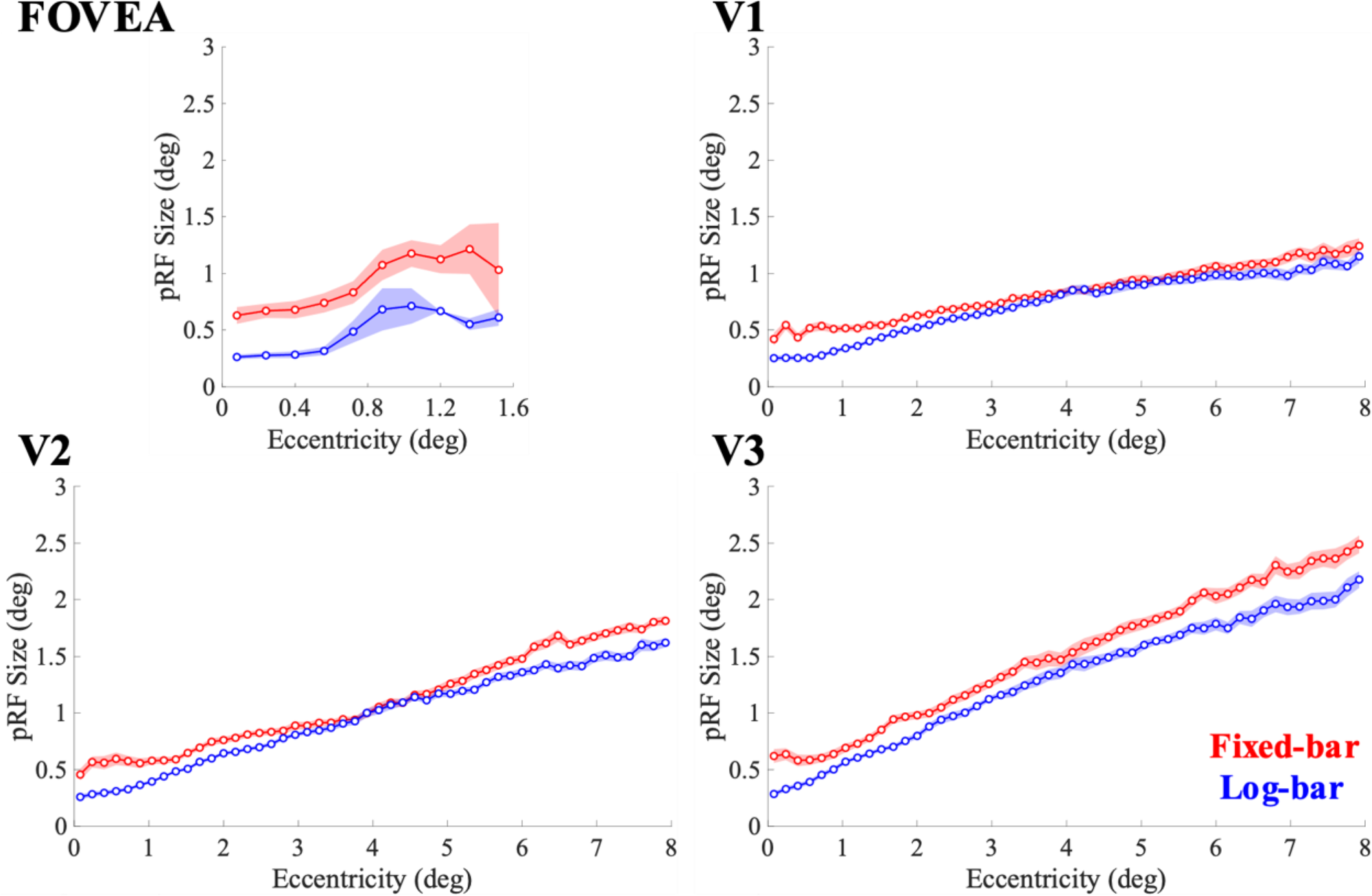
pRF size estimates binned as a function of eccentricity. Solid lines represent the average across participants for the fixed-bar (red) and log-bar (blue) stimuli and the shaded area represents the ±1 standard error of the mean.

Coefficients for slope and intercept were acquired by fitting a regression line to eccentricity and pRF size estimates. There was a main effect of stimulus type (*F*(1,164) = 120.14, *p* < 0.0001, η^2^=0.42) and ROI (*F*(3,164) = 5.83, *p* < 0.0001, η^2^=0.10) on the intercept of pRF size by eccentricity. The intercept parameter was larger for the fixed-bar stimulus than the log-bar stimulus. The smallest intercept values were from V1 and V2, followed by V3, and then the fovea. There was an interaction effect of stimulus type by ROI (*F*(3,164) = 5.50, *p* = 0.0013, η^2^=0.09) where the fixed-bar stimulus had larger intercepts than the log-bar stimulus, especially in the foveal confluence. There was no difference in slope coefficient for any variables examined.

### pRF Model Simulations

**Figure 7** compares the ability to recover accurate pRF estimates from fixed and log-bar stimuli. As shown in **Panel A**, consistent with our data in Figure 5, the fixed-bar stimulus overestimates pRF sizes at eccentricities of less than 1.5 degrees. **Panel B** plots these same data as a function of the original pRF sizes used to generate the simulations. Recovered pRF sizes from the log-bar stimuli closely matched the simulation pRF sizes across all eccentricities. In contrast, estimates made using fixed-bar stimulus systematically overestimated pRF sizes when the simulated initial pRF values were < 0.2°, which in our model represented regions of less than 1.5° eccentricity. As described in the introduction, the size of these biases is determined by the mismatch between the size of the pRF and the width and speed of the traveling bar.

**Figure 7.**
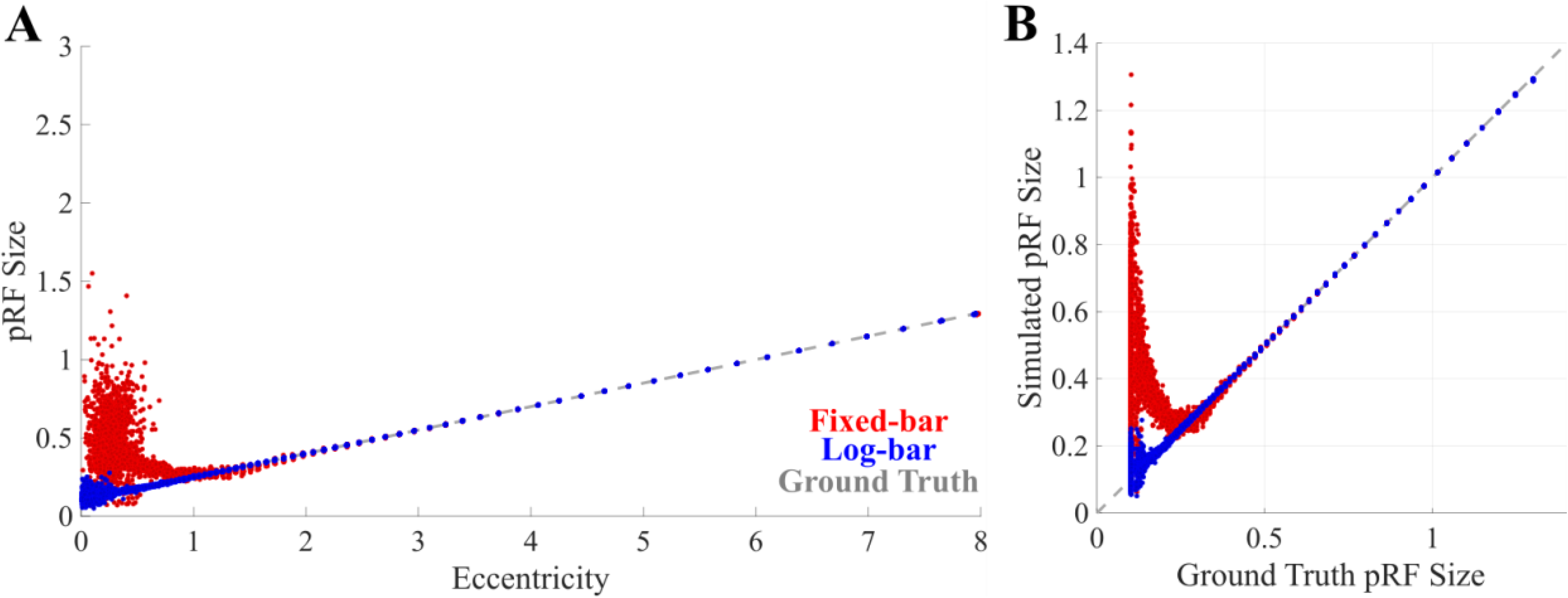
Simulated pRF size estimates and comparisons to ground truth. (A) Simulated pRF size as a function of eccentricity. (B) Simulated pRF size vs. ground truth pRF size. The colors indicate pRF estimates from the fixed-bar (red), log-bar (blue), and ground truth (dashed gray).

## Discussion

This study shows how pRF estimates can be improved by using a logarithmically-distorted drifting bar stimulus instead of the standard fixed-bar stimulus. We found that across early visual areas, the log-bar stimulus (1) produced more reliable pRF estimates of polar angle, eccentricity, and size; (2) produced more pRFs near the foveal representation; and (3) produced pRFs that were much smaller in size and closer to the fovea (more closely resembling the what is believed to be the underlying ‘ground-truth’) than those produced with the traditional drifting bar stimulus.

The limitations of the current study arise from technical capabilities and design choices. First, we were unable to stimulate the visual field beyond 8° eccentricity due to equipment constraints, and we chose to distort the log-bar with only a single factor (*k* = 5) to approximately match CMF properties of human early visual cortex. Therefore, we are unable to consider the effects the log-bar stimulus would have in more eccentric visual locations and visual areas beyond early visual cortex (i.e., V4 – V7). It is possible that our distortion factor would not generalize to extrastriate visual cortex, and these higher-order visual areas may require their own optimized stimulus configuration. Second, we used a 100% contrast flickering checkerboard as the background texture for our stimuli. Flickering checkerboard stimuli are known to strongly drive neural activity; however, it may not be the optimal background texture. Visual cortex has demonstrated tuning for 1/f spectrum noise (Isherwood et al., 2017) and the Human Connectome Project implemented their stimulus with 1/f noise with embedded objects (Benson et al., 2018). It is possible that a combination of an optimized background texture and optimized stimulus aperture, like our log-bar stimulus, could lead to better pRF estimates and model performance, especially in higher-order visual areas. Finally, our pRF model did not incorporate non-linear visual computations like compressive spatial (Kay et al., 2013) or temporal (Zhou et al., 2018) summation, surround suppression (Zuiderbaan et al., 2012), or normalization (Aqil et al., 2021). In the spatial domain, incorporating these nonlinearities tends to reduce pRF size estimates, and linear model can overestimate pRF size when the underlying system exhibits nonlinear, subadditive behavior (Kay et al., 2013). However, these models tend to find that V1 is approximately linear, and non-linearities increase across the visual hierarchy. In contrast, our log-bar stimulus had the strongest influences on pRFs in the foveal confluence, suggesting that both a log-bar stimulus and a model capable of capturing spatial and temporal non-linearities will be needed to accurately estimate pRFs.

In summary, a logarithmically-distorted drifting bar stimulus, locally optimized to match the receptive fields of the corresponding region of cortical space, produces more accurate and reliable pRF estimates than a standard drifting bar stimulus, especially for pRFs located near central vision.

## Acknowledgments

This research was supported by grants from NIH National Eye Institute Grant R01EY014645 (IF) and the University of Washington Center for Human Neuroscience (CHN).

The data and code reported in this article are available in the OpenNeuro repository (https://doi.org/10.18112/openneuro.ds004698.v1.0.1).

## References

Alvarez, I., De Haas, B. A., Clark, C. A., Rees, G., & Schwarzkopf, D. S. (2015). Comparing different stimulus configurations for population receptive field mapping in human fMRI. Frontiers in Human Neuroscience, 9. 10.3389/fnhum.2015.00096

Andersson, J. L. R., Skare, S., & Ashburner, J. (2003). How to correct susceptibility distortions in spin-echo echo-planar images: Application to diffusion tensor imaging. NeuroImage, 20(2), 870–888. 10.1016/S1053-8119(03)00336-7

Aqil, M., Knapen, T., & Dumoulin, S. O. (2021). Divisive normalization unifies disparate response signatures throughout the human visual hierarchy. Proceedings of the National Academy of Sciences, 118(46), e2108713118. 10.1073/pnas.2108713118

Barbot, A., Das, A., Melnick, M. D., Cavanaugh, M. R., Merriam, E. P., Heeger, D. J., & Huxlin, K. R. (2021). Spared perilesional V1 activity underlies training-induced recovery of luminance detection sensitivity in cortically-blind patients. Nature Communications, 12(1), 6102. 10.1038/s41467-021-26345-1

Baseler, H. A., Gouws, A., Haak, K. V., Racey, C., Crossland, M. D., Tufail, A., Rubin, G. S., Cornelissen, F. W., & Morland, A. B. (2011). Large-scale remapping of visual cortex is absent in adult humans with macular degeneration. Nature Neuroscience, 14(5), 649–655. 10.1038/nn.2793

Behzadi, Y., Restom, K., Liau, J., & Liu, T. T. (2007). A component based noise correction method (CompCor) for BOLD and perfusion based fMRI. NeuroImage, 37(1), 90–101. 10.1016/j.neuroimage.2007.04.042

Benson, N. C., Butt, O. H., Datta, R., Radoeva, P. D., Brainard, D. H., & Aguirre, G. K. (2012). The Retinotopic Organization of Striate Cortex Is Well Predicted by Surface Topology. Current Biology, 22(21), 2081–2085. 10.1016/j.cub.2012.09.014

Benson, N. C., Jamison, K. W., Arcaro, M. J., Vu, A. T., Glasser, M. F., Coalson, T. S., Van Essen, D. C., Yacoub, E., Ugurbil, K., Winawer, J., & Kay, K. (2018). The Human Connectome Project 7 Tesla retinotopy dataset: Description and population receptive field analysis. Journal of Vision, 18(13), 23–23. 10.1167/18.13.23

Benson, N. C., & Winawer, J. (2018). Bayesian analysis of retinotopic maps. eLife, 7, e40224. 10.7554/eLife.40224

Binda, P., Thomas, J. M., Boynton, G. M., & Fine, I. (2013). Minimizing biases in estimating the reorganization of human visual areas with BOLD retinotopic mapping. Journal of Vision, 13(7), 13. 10.1167/13.7.13

Brainard, D. H. (1997). The Psychophysics Toolbox. Spatial Vision, 10(4). 10.1163/156856897X00357

Brewer, A. A., & Barton, B. (2012). Effects of healthy aging on human primary visual cortex. Health, 4(9), Article 9. 10.4236/health.2012.429109

Cox, R. W., & Hyde, J. S. (1997). Software tools for analysis and visualization of fMRI data. NMR in Biomedicine, 10(4–5), 171–178. 10.1002/(SICI)1099-1492(199706/08)10:4/5<171::AID-NBM453>3.0.CO;2-L

Dale, A. M., Fischl, B., & Sereno, M. I. (1999). Cortical Surface-Based Analysis: I. Segmentation and Surface Reconstruction. NeuroImage, 9(2), 179–194. 10.1006/nimg.1998.0395

Daniel, P. M., & Whitteridge, D. (1961). The representation of the visual field on the cerebral cortex in monkeys. The Journal of Physiology, 159(2), 203–221. 10.1113/jphysiol.1961.sp006803

Dumoulin, S. O., & Wandell, B. A. (2008). Population receptive field estimates in human visual cortex. NeuroImage, 39(2), 647–660. 10.1016/j.neuroimage.2007.09.034

Duncan, R. O., & Boynton, G. M. (2003). Cortical Magnification within Human Primary Visual Cortex Correlates with Acuity Thresholds. Neuron, 38(4), 659–671. 10.1016/S0896-6273(03)00265-4

Esteban, O., Markiewicz, C. J., Blair, R. W., Moodie, C. A., Isik, A. I., Erramuzpe, A., Kent, J. D., Goncalves, M., DuPre, E., Snyder, M., Oya, H., Ghosh, S. S., Wright, J., Durnez, J., Poldrack, R. A., & Gorgolewski, K. J. (2019). fMRIPrep: A robust preprocessing pipeline for functional MRI. Nature Methods, 16(1), 111–116. 10.1038/s41592-018-0235-4

Greve, D. N., & Fischl, B. (2009). Accurate and robust brain image alignment using boundary-based registration. NeuroImage, 48(1), 63–72. 10.1016/j.neuroimage.2009.06.060

Haak, K. V., Cornelissen, F. W., & Morland, A. B. (2012). Population Receptive Field Dynamics in Human Visual Cortex. PLOS ONE, 7(5), e37686. 10.1371/journal.pone.0037686

Harvey, B. M., & Dumoulin, S. O. (2011). The Relationship between Cortical Magnification Factor and Population Receptive Field Size in Human Visual Cortex: Constancies in Cortical Architecture. Journal of Neuroscience, 31(38), 13604–13612. 10.1523/JNEUROSCI.2572-11.2011

Himmelberg, M. M., Kurzawski, J. W., Benson, N. C., Pelli, D. G., Carrasco, M., & Winawer, J. (2021). Cross-dataset reproducibility of human retinotopic maps. NeuroImage, 244, 118609. 10.1016/j.neuroimage.2021.118609

Huber, E., Chang, K., Alvarez, I., Hundle, A., Bridge, H., & Fine, I. (2019). Early Blindness Shapes Cortical Representations of Auditory Frequency within Auditory Cortex. Journal of Neuroscience, 39(26), 5143–5152. 10.1523/JNEUROSCI.2896-18.2019

Infanti, E., & Schwarzkopf, D. S. (2020). Mapping sequences can bias population receptive field estimates. NeuroImage, 211, 116636. 10.1016/j.neuroimage.2020.116636

Isherwood, Z. J., Schira, M. M., & Spehar, B. (2017). The tuning of human visual cortex to variations in the 1/fα amplitude spectra and fractal properties of synthetic noise images. NeuroImage, 146, 642–657. 10.1016/j.neuroimage.2016.10.013

Kay, K. N., Winawer, J., Mezer, A., & Wandell, B. A. (2013). Compressive spatial summation in human visual cortex. Journal of Neurophysiology, 110(2), 481–494. 10.1152/jn.00105.2013

Lage-Castellanos, A., Valente, G., Senden, M., & De Martino, F. (2020). Investigating the Reliability of Population Receptive Field Size Estimates Using fMRI. Frontiers in Neuroscience, 14. 10.3389/fnins.2020.00825

Linhardt, D., Pawloff, M., Hummer, A., Woletz, M., Tik, M., Ritter, M., Schmidt-Erfurth, U., & Windischberger, C. (2021). Combining stimulus types for improved coverage in population receptive field mapping. NeuroImage, 238, 118240. 10.1016/j.neuroimage.2021.118240

Pelli, D. G. (1997). The VideoToolbox software for visual psychophysics: Transforming numbers into movies. Spatial Vision, 10(4), 437–442. 10.1163/156856897X00366

Satterthwaite, T. D., Elliott, M. A., Gerraty, R. T., Ruparel, K., Loughead, J., Calkins, M. E., Eickhoff, S. B., Hakonarson, H., Gur, R. C., Gur, R. E., & Wolf, D. H. (2013). An improved framework for confound regression and filtering for control of motion artifact in the preprocessing of resting-state functional connectivity data. NeuroImage, 64, 240–256. 10.1016/j.neuroimage.2012.08.052

Schira, M. M., Wade, A. R., & Tyler, C. W. (2007). Two-Dimensional Mapping of the Central and Parafoveal Visual Field to Human Visual Cortex. Journal of Neurophysiology, 97(6), 4284–4295. 10.1152/jn.00972.2006

Schwartz, E. L. (1980). Computational anatomy and functional architecture of striate cortex: A spatial mapping approach to perceptual coding. Vision Research, 20(8), 645–669. 10.1016/0042-6989(80)90090-5

Schwarzkopf, D. S., Anderson, E. J., Haas, B. de, White, S. J., & Rees, G. (2014). Larger Extrastriate Population Receptive Fields in Autism Spectrum Disorders. Journal of Neuroscience, 34(7), 2713–2724. 10.1523/JNEUROSCI.4416-13.2014

Senden, M., Reithler, J., Gijsen, S., & Goebel, R. (2014). Evaluating Population Receptive Field Estimation Frameworks in Terms of Robustness and Reproducibility. PLOS ONE, 9(12), e114054. 10.1371/journal.pone.0114054

Tisdall, M. D., Hess, A. T., Reuter, M., Meintjes, E. M., Fischl, B., & van der Kouwe, A. J. W. (2012). Volumetric navigators for prospective motion correction and selective reacquisition in neuroanatomical MRI. Magnetic Resonance in Medicine, 68(2), 389–399. 10.1002/mrm.23228

Tustison, N. J., Avants, B. B., Cook, P. A., Zheng, Y., Egan, A., Yushkevich, P. A., & Gee, J. C. (2010). N4ITK: Improved N3 Bias Correction. IEEE Transactions on Medical Imaging, 29(6), 1310–1320. 10.1109/TMI.2010.2046908

van Dijk, J. A., de Haas, B., Moutsiana, C., & Schwarzkopf, D. S. (2016). Intersession reliability of population receptive field estimates. NeuroImage, 143, 293–303. 10.1016/j.neuroimage.2016.09.013

Vanni, S., Henriksson, L., & James, A. C. (2005). Multifocal fMRI mapping of visual cortical areas. NeuroImage, 27(1), 95–105. 10.1016/j.neuroimage.2005.01.046

Zeidman, P., Silson, E. H., Schwarzkopf, D. S., Baker, C. I., & Penny, W. (2018). Bayesian population receptive field modelling. NeuroImage, 180, 173–187. 10.1016/j.neuroimage.2017.09.008

Zhang, Y., Brady, M., & Smith, S. (2001). Segmentation of brain MR images through a hidden Markov random field model and the expectation-maximization algorithm. IEEE Transactions on Medical Imaging, 20(1), 45–57. 10.1109/42.906424

Zhou, J., Benson, N. C., Kay, K. N., & Winawer, J. (2018). Compressive Temporal Summation in Human Visual Cortex. Journal of Neuroscience, 38(3), 691–709. 10.1523/JNEUROSCI.1724-17.2017

Zuiderbaan, W., Harvey, B. M., & Dumoulin, S. O. (2012). Modeling center–surround configurations in population receptive fields using fMRI. Journal of Vision, 12(3), 10. 10.1167/12.3.10

